# Establishment and characterisation of primary skeletal muscle cell cultures from patients with advanced Chronic Kidney Disease

**DOI:** 10.1101/2020.11.16.384263

**Authors:** Luke A Baker, Tom F O’Sullivan, Kate A Robinson, Zoe Redshaw, Matthew Graham-Brown, Robert U Ashford, Alice C Smith, Andrew Philp, Emma L Watson

## Abstract

Skeletal muscle wasting and dysfunction is a common characteristic of non-dialysis dependent chronic kidney disease (NDD-CKD). The mechanisms by which this occurs are not clearly understood and one reason for this is a lack of well controlled *in-vitro* methodologies to simulate NDD-CKD induced muscle wasting for mechanistic investigation at the cellular level. Here we sought to conduct the initial investigations into developing a CKD-induced skeletal muscle model for use as a mechanistic analysis tool as well as a test bed for potential novel therapeutics in this population. Human derived muscle cells (HDMCs) were isolated from n=5 NDD-CKD patients and n=3 matched healthy controls (HC) and taken through proliferation and differentiation phases in cell culture. Upon comparison of the 2 donor types, significantly greater mRNA expression of myogenic markers was noted in the NDD-CKD cultures in comparison to HC cultures, which was carried through to greater mRNA expression of myosin heavy chains (MyHCs) post differentiation. However, this was not carried over to protein expression where Pax7 and MyoD were seen to be expressed to a greater extent in HC cultures. mRNA expression markers of protein degradation were noted to be elevated in NDD-CKD cultures in comparison to HC cultures. In light of our findings, future work should seek to investigate the role of the ‘CKD environment’ as well as mechanisms implicated in transcription regulation to further advance the current model development as well as the mechanistic understanding of skeletal muscle wasting in CKD.

## Introduction

Skeletal muscle wasting and dysfunction is a common characteristic of chronic kidney disease (CKD) (21). This can potentially limit physical activity, resulting in a downward spiral of atrophy, deconditioning, disuse and poor outcomes (19, 20). These are important clinical problems as they negatively impact upon quality of life and are associated with increased rates of morbidity and mortality as well as contributing to the continued rise of health and social care costs worldwide (10). The mechanisms underlying muscle wasting and dysfunction are not well explored in human CKD. To date, this important clinical question has been investigated using either animal models of CKD (8), which do not always replicate what we see in our CKD patients, or muscle biopsies(3, 6, 16, 19, 20), which represent a single time point and are limited with regard to mechanistic investigation. The development of a biologically relevant model of human skeletal muscle culture model is therefore important for the understanding of muscle loss and dysfunction in human CKD.

Skeletal muscle models of atrophy are not uncommon, with recent work systematically defining a high through put test bed for the screening of therapeutic interventions (2). However, studies to this point seeking to model the characteristics of CKD skeletal muscle *in-vitro* have looked to manipulate the proposed systemic environment of CKD and expose this environment to immortalised cells lines. Though this has proven effective (8), it fails to represent the phenotypic adaptation of CKD muscle, characteristics that could only be replicated through the use of tissue derived from the patients themselves. Human derived muscle cells (HDMCs) have been shown to retain their phenotypic traits well, including disturbances in metabolic processes(1, 12). Therefore, we sought to develop a primary culture model established from muscle biopsies collected from patients with advanced non-dialysis dependent CKD (NDD-CKD) in which more rigorous experimentation can be performed. Skeletal muscle primary cells are increasingly used to research muscle disorders which is a feature of other chronic illnesses (9, 14), but are yet to be utilised to investigate muscle wasting in human CKD. Such a model would provide an opportunity for mechanistic investigations specific to this disease state, enabling progression of the current knowledge base and the development of CKD specific therapeutic interventions.

## Methods

### Sample collection

A single muscle biopsy (approximately 50mg wet weight) was collected from n = 5 NDD-CKD patients (CKD stage 3b-4) using the micro biopsy technique after an overnight fast, as described previously (6). Exclusion criteria included uncontrolled diabetes (HbA1c > 9%), cancer, inflammatory and musculoskeletal disorders. Muscle biopsies were also collected from n = 3 age and sex matched healthy controls (HC) with no significant medical history, using an open biopsy technique as these individuals were already under going planned minor orthopaedic surgery. After dissection of any visible fat and connective tissue, samples were placed in 5mL Hams F10 media (Gibco) on ice for two hours prior to satellite cell isolation. Ethical approval was given by the National Research Ethics Committee (Ref:15/EM/0467). All patients gave written informed consent and the trial was conducted in accordance with the Declaration of Helsinki. This study is registered with the ISRCTN (Ref: 18221837).

### Satellite cell isolation procedure

Muscle tissue was washed three times in HamsF10 (containing 1% penicillin streptomycin and 1% Gentamycin), minced into small fragments and enzymatically digested in two incubations with collagenase IV (1mg/mL), BSA (5mg/mL) and trypsin (500μl/mL) at 37°C with gentle agitation. The resultant supernatant was added to FBS on ice, strained through a 70μm nylon filter and centrifuged at 800 g for 7 minutes. The cells were washed in Hams F10 with 1% penicillin streptomycin and pre-plated on uncoated 9cm^2^ petris in 3mL growth medium (GM; Hams F10 Glutamax, 20% FBS, 1% penicillin streptomycin, 1% fungazone) for 3 h. The cell suspension was then moved to collagen I coated 25 cm^2^ flasks and kept in standard culture environmental conditions (37°C, 5 % CO_2_). For the expansion of satellite cell populations, cells were grown to approximately 70% confluence in GM that was changed every other day. Cells were subsequently trypsinised (Sigma-Aldrich) and counted using the trypan blue exclusion method.

### Cell Culture

Myoblasts were seeded on to collagen coated (Sigma) 6-well plates at 100,000 cells/well and cultured to 70% confluence. At this point cells were then switched to differentiation medium (DM; DMEM, 2% horse serum & 1% penicillin streptomycin) for 7 days for the induction of myotube formation. Cultures were harvested at 70 % confluence (0D), 3 days (3D) and 7 days (7D) post DM for analysis of markers of proliferation, myogenesis, maturity and degradation signalling.

### Quantitative RT-PCR

Total RNA was extracted from cells using 1mL/well Trizol and 1μg RNA was reverse transcribed to cDNA using an AMV reverse transcription system (Promega, Madison, WI, USA). Primers, probes and internal controls for all genes were supplied as TaqMan™ gene expression assays: Myogenin - Hs01072232_m1; MyoD - Hs02330075_g1; Myf-5 - Hs00929416_g1; ki-67 - Hs04260396_g1; Pax-7 - Hs00242962_m1; MyHC-1 - Mm01332489_m1; MyHC-2 - Hs00430042_m1; MyHC-3 - Hs01074230_m1; MyHC-7 - Mm00600555_m1; MyHC-8 - Hs00267293_m1; Fbxo32 - Hs01041408_m1; TRIM63 – Hs00822397_m1 and TBP - Hs00427620_m1 was used as a reference gene. All reactions were carried out in a 20 μl volume: 1 μl cDNA, 10 μl 2X Taqman Mastermix, 8 μl water, 1 μl primer/probe on an Agilent Biosystem Light Cycler with the following conditions, 95°C 15 s, followed by 40x at 95°C for 15 s and 60°C for 1 min. The Ct values from the target gene were normalized to TBP and expression levels calculated according to 2−Δ Ct method to determine fold changes.

### Western Blotting

Total protein was extracted from the phenol phase of the Trizol extraction using the guanine method. Protein concentration was determined using the Bio-Rad DC Protein Assay. Lysates containing 30 μg protein were subjected to SDS-PAGE using 10% Criterion TGX Stain-Free gels (BioRad, UK) on a mini-Protean Tetra system (Bio-Rad, UK). Stain-free imaging was performed using a ChemiDoc MP imager (Bio-Rad) with a 5-minute UV activation step prior to transfer. Proteins were transferred onto nitrocellulose membranes, blocked for 1 h with Tris-buffered saline with 5% (w/v) skimmed milk and 0.1% (v/v) Tween-20 detergent. Membranes were incubated with the primary antibody overnight as follows: Pax-7 (Abcam, 1:1000); MyoD (Abcam, 1:750); Myogenin (ThermoFisher, 1:1000). Horseradish Peroxidase (HRP) linked secondary anti-mouse/rabbit secondary antibodies (Dako, Aglient, UK) were used at 1:1500 for 2h at room temperature. Blots were visualised using ECL Reagents (Geneflow, UK) captured using a ChemiDoc MP imager (Bio-Rad). Total protein was normalised to the stain free blot image providing quantification of total protein.

### Statistical Analysis

Data is presented as mean ± standard deviation unless otherwise stated. All data was tested for normality using the Shapiro-Wilk test. Non-normally distributed data was either log-transformed prior to analysis or a non-parametric equivalent was used as appropriate. To determine the effect of condition over time and between groups of donors a two-way repeated measures ANOVA was used (time x condition) with Bonferroni post-hoc analysis used for identification of where significance lay within the data set. Statistical analysis was carried out using IBM SPSS 25 software (IBM, Chicago, IL). Statistical significance was accepted as p ≤ 0.05, with the amplitude of significance indicated throughout.

## Results

No significant differences were noted between the two groups regarding gender (HC 1M:2F; NDD-CKD 2M:3F) or age (HC 58.66 ± 14.74 vs. NDD-CKD 54.53 ± 15.53 yr) with significant differences only being noted in eGFR (HC 75 ± 5.56 vs. NDD-CKD 22.25 ± 13.22 mL/min/1.73 m^2^, p ≤ 0.001).

Gene expression analysis revealed the expected time-course patterns, with proliferative markers (Ki-67, Myf-5 & Pax7) peaking at the point of confluence (D0), followed by increases in markers of myogenesis (MyoD & Myogenin) at D3 and subsequent timely increases in MyHC expression as cultures progressed to D7. Across all markers, differences between cell culture passages 1-4 were seen to be non-significant (p > 0.05), thus all of the data displayed includes combined passages. Upon comparison of the two donor groups, significantly greater mRNA expression of Pax-7 (p = 0.005), Myf-5 (p = 0.0004) and MyoD (p = 0.0.001) was noted in CKD-derived cultures at D0, with higher expression also observed at D3 for Pax7 (p=0.055) and MyoD (p = 0.017), compared to HC cultures. Further to this, at D7, significantly greater expression of all MyHC’s was observed in CKD-derived cultures compared to HC-derived cultures (MyHC-1 p = 0.014; MyHC-2 p = 0.003; MyHC-3 p < 0.0001; MyHC-7 p = 0.022; MyHC-8 p = 0.005).

Considering that upregulated E3 ubiquitin ligase signalling has been strongly implicated in CKD-related muscle wasting as key regulators of the ubquitin proteosome system (UPS)(4), we sought to investigate the extent to which this phenotype was preserved in culture. Time-dependent increases in MafBX (encoded by the Fbxo32 gene) and Murf-1 (encoded by the TRIM63 gene) were observed in both donor cultures, with significantly greater expression of Fbxo32 detected at D7 in CKD versus HC cultures (p = 0.0001). No significant differences were noted in TRIM63 expression between donors at D7 (p = 0.17).

**Figure 1:**
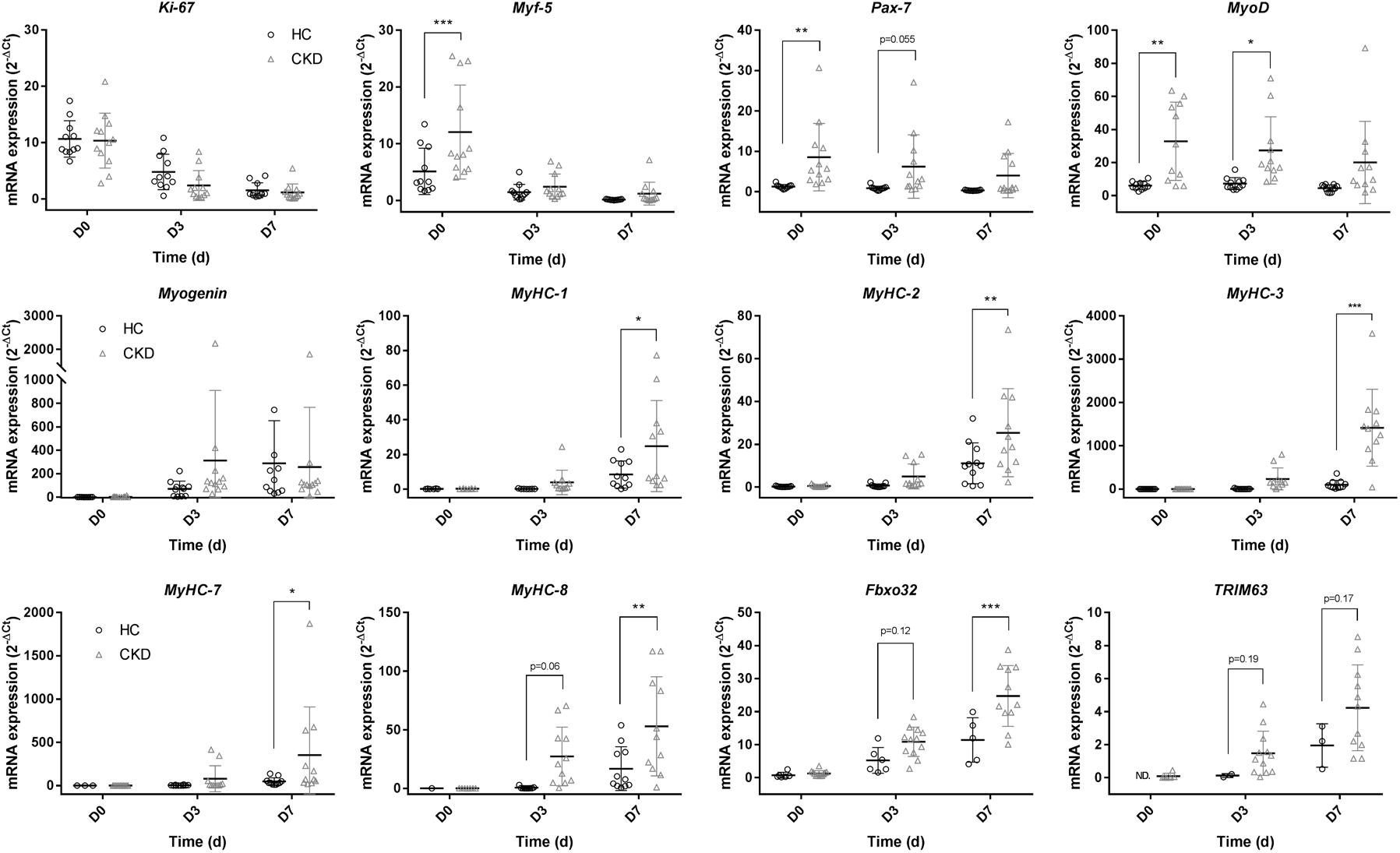
mRNA gene expression comparison between HC and NDD-CKD cultures over a 7 day time course. mRNA gene expression analysis comparing HC and NDD-CKD at days 0, 3 and 7 for quantification of markers of: Proliferation: ki-67, Myf-5 & Pax-7; Myogenic factors: MyoD & Myogenin; Maturity: MyHC-1, MyHC-2, MyHC-3, MyHC-7 & MyHC-8; Protein breakdown: Fbxo32 & TRIM63. Data presented as means ± SD. * indicating p ≤ 0.05, ** indicating p ≤ 0.01, *** indicating p ≤ 0.001.

To provide context for the expression analysis, western blot analysis was conducted to semi quantify the functional protein levels to see if changes at the gene expression level are carried over into translational changes for the myogenic markers MyoD and Myogenin at each time point. In contrast to the mRNA data, Pax-7 protein expression was seen to be significantly higher in HC cultures in comparison to CKD cultures at D0 (p = 0.0058). Trends to greater levels of MyoD were also noted in HC over CKD cultures at the 3D time point (p = 0.08), again reversing that seen in gene expression data. No differences were noted across the time points between the two donors cultures in Myogenin (p > 0.05).

**Figure 2:**
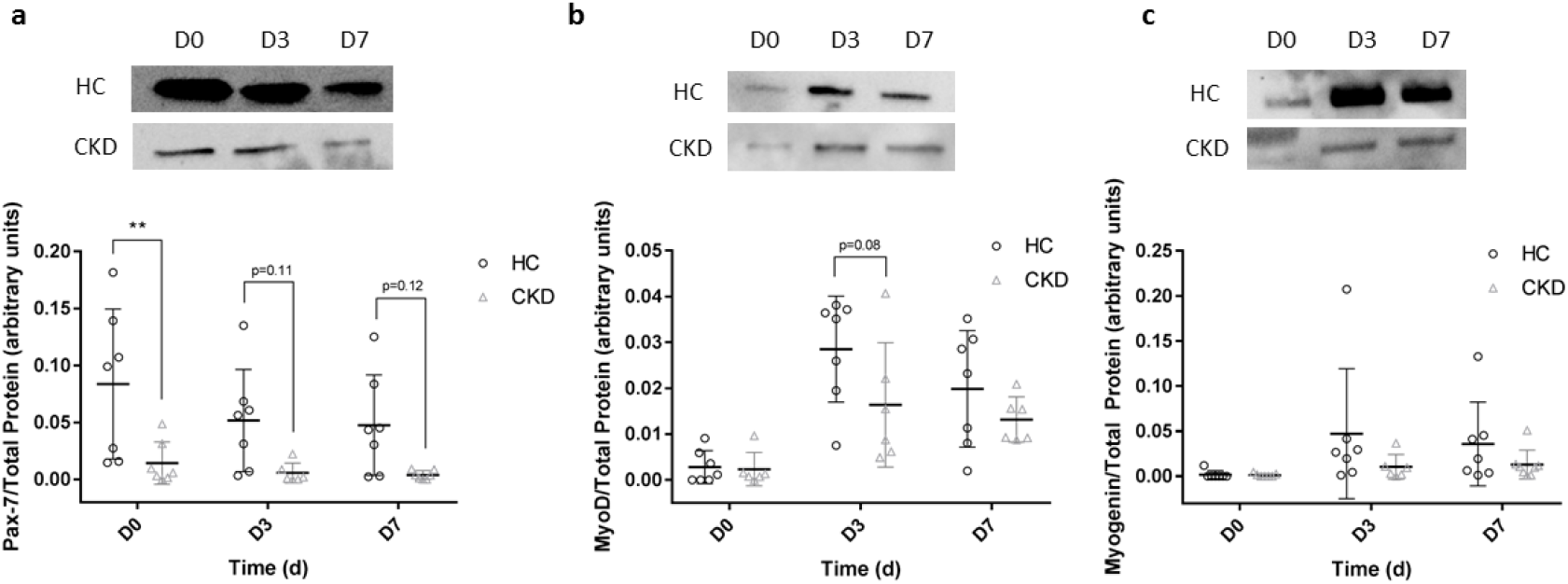
Protein expression comparison between HC and NDD-CKD cultures over a 7 time course. Protein expression analysis comparing HC and NDD-CKD cultures at days 0, 3 and 7: a) Pax-7 relative to total protein load; b) MyoD relative to total protein load; c) Myogenin relative to total protein load. Data presented as means ± SD. ** indicating p ≤ 0.01.

## Discussion

The work presented, to the authors knowledge, is the first to attempt to replicate the muscle wasting phenotype observed in NDD-CKD in an *in-vitro* setting for mechanistic experimentation. The findings were both suprising and of great interest, providing many lines of enquiry for future investgations. Expression of Ki-67, Pax-7 and Myf-5 have been used throughout the literature to indicate increased rates of proliferation(7). In the current investgation we were able to show that HDMC from NDD-CKD have greater mRNA expression levels of Pax-7 and Myf-5 in comparison to those derived from matched HC. A possible explanation for this could be that NDD-CKD pateints are systemically inflammed(17) and thus cells derived from NDD-CKD patients have been predisposed to an inflammatory environment. It has been suggested that certain inflammatory cytokines (eg. IL-6) may have a homeostatic role to induce proliferation in order for the repair and regeneration of muscle fibres during injury or infection(13). Previous literature has demonstrated ‘muscle memory’ of myoblasts *in-vitro* (15) in response to TNF-α exposure, and suggested an ‘epi-memory’ of HDMCs when cells are analysed from healthy donors during ageing. Consequently, previous exposure to increased IL-6 could feasibly result in epigenetic adaptation to the systemic environment, potentially leading to enhanced proliferation *in vitro*, thus eluding to an epigenetic adaptation of skeletal muscle to the systemic NDD-CKD enviroment. If further investgations are able to confirm the proposed hypothesis and its underpinning mechanisms, this would not only aid our development of an *in-vitro* model of CKD related wasting, but also broaden our understanding of the mechanisms contributing to this phenotype, thus highlighting opportunities for therapeutic intervention.

Upon analysis of the mRNA expression of an array fo MyHC’s, CKD cultures showed a general upregulation in comparison to HC cultures. This was coupled with greater mRNA expression of both MafBX and Murf-1 in CKD cultures also, or trends to reductions where biological repeats were limited. Regarding the MyHC’s, these preliminary findings are the opposite to what would be expected but provide an interesting biological insight. We speculatively propose that enhanced expression of MyHC in CKD cultures was enhanced to compensate for the increased E3-ligase expression carried over from the CKD inflammatory enviroment the cells were exposed to *in-situ*. Our work presents an opportunity for further mechanistic investgations of the propsed hypothosies to further our knowleadge base regarding muscle wasting in NDD-CKD patient populations.

With proliferative markers being noted to be higher at the gene expression level, protein analysis was conducted to investgate if mRNA transcription correlated to translation. Both Pax-7 and MyoD saw findings reversed compared to the mRNA data, with HC cultures having greater protein expression compared to CKD cultures. Findings at the protein level replicate more closely those previously described *in-vivo* making such a finding important for the progression of the current model. From a mechanistic stand point, this finding suggests that CKD derived cells potentially have reduced capacity to synthesise proteins, suggesting either a dysregulation of myogenic related micro-RNAs or ribosomal biogenesis dysfunction. The dysregulation of micro-RNAs involved in myogenic regulation has been reported previously in animal models(8, 11), but investgations in human CKD are still lacking and warrant greater attention. To the authors knowleadge, investgations into ribosomal biogenesis dysfunction in human CKD is still an emerging area of research(18). Previous work has shown the role of such processes in the develoment of skeletal muscle wasting (5) but this has yet to be applied to our pateint population.

In conclusion, our novel investgations presented here depict the properties of HDMCs derived from NDD-CKD compared to matched controls. We have attempted to create an *in-vitro* model of CKD muscle wasting, in order to conduct further mechanistic investgations for patient benefit. Future work should seek to investgate the role of the ‘NDD-CKD enviroment’ using this methodology to assertain its contribution to wasting in this population. Further to this, in depth investgations into the role of micro-RNAs and ribosome biogenesis in muscle wasting should be conducted in this population in response to the the current findings.

